# Influence of multiple hypothesis testing on reproducibility in neuroimaging research

**DOI:** 10.1101/488353

**Authors:** Tuomas Puoliväli, Satu Palva, J. Matias Palva

## Abstract

**Background:** Reproducibility of research findings has been recently questioned in many fields of science, including psychology and neurosciences. One factor influencing reproducibility is the simultaneous testing of multiple hypotheses, which increases the number of false positive findings unless the p-values are carefully corrected. While this multiple testing problem is well known and has been studied for decades, it continues to be both a theoretical and practical problem.

**New Method:** Here we assess the reproducibility of research involving multiple-testing corrected for family-wise error rate (FWER) or false discovery rate (FDR) by techniques based on random field theory (RFT), cluster-mass based permutation testing, adaptive FDR, and several classical methods. We also investigate the performance of these methods under two different models.

**Results:** We found that permutation testing is the most powerful method among the considered approaches to multiple testing, and that grouping hypotheses based on prior knowledge can improve power. We also found that emphasizing primary and follow-up studies equally produced most reproducible outcomes.

**Comparison with Existing Method(s):** We have extended the use of two-group and separate-classes models for analyzing reproducibility and provide a new open-source software “MultiPy” for multiple hypothesis testing.

**Conclusions:** Our results suggest that performing strict corrections for multiple testing is not sufficient to improve reproducibility of neuroimaging experiments. The methods are freely available as a Python toolkit “MultiPy” and we aim this study to help in improving statistical data analysis practices and to assist in conducting power and reproducibility analyses for new experiments.

## 1. Introduction

The reproducibility of published research has been recently called into question in many fields, including psychology and neurosciences (Button et al., 2013; Open Science Collaboration, 2015; Baker, 2016; Poldrack et al., 2017; Poldrack, 2019). Reproducibility is affected by many factors, one of which is the simultaneous testing of multiple hypotheses (Ioannidis, 2005), which increases the number of false positive findings unless the corresponding p-values are appropriately corrected. Hence, there is demand for tools to evaluate multiple-testing data-analysis plans and to perform the required corrections using appropriate methods. Although numerous techniques exist for multiple hypothesis testing, it has remained incompletely understood how much the choice of method and relative emphasis on the primary study over follow-up study, or vice versa, influences the reproducibility of the observations. Here we developed a Python-based software library that implements techniques for controlling the family-wise error rate (FWER) and the false discovery rate (FDR) and then developed a novel model-based approach to compare their relative performance in terms of their power, false positive rate, and reproducibility.

We assess the differences between the methods that control the FWER and FDR by comparing their performance using simulated data generated under two related spatial models. The first one is an extension of the classic two-group model that consists of distinct signal and noise regions (Bennett et al., 2009). The second one is an extension of Efron’s (2008) separate-classes model; it combines two two-group models for being able to represent distributed effects. These models can be used to represent hypothetical effects in neurophysiological and neuroimaging data such as evoked or induced activity in an electroencephalography (EEG) or magnetoencephalography (MEG) time-frequency analysis, or relatively focal effects in functional or anatomical magnetic resonance imaging (MRI) data. In addition, these models allow performing numerical prospective power analyses to facilitate planning of new experiments, including the determination of sample and effect sizes that are required for observing true effects reproducibly. Since fundamental effects in systems-level neuroscience data are often spatially or temporally continuous (Penny & Friston, 2003; Heller et al., 2006; Chumbley et al., 2010), as well as distributed, these model are suitable in analyzing a wide range research questions. Moreover, only few previous studies to date have investigated properties of multiple testing procedures under models with two or more simultaneous distinct effects. Efron (2008) analyzed inference under a separate-classes model, which was motivated by diffusion tensor imaging data from dyslexic and control participants indicating distinct effects in anterior and posterior parts of the brain. Their results showed that since the underlying data had distinct effects and structures, performing two separate corrections was better than a single combined analysis (Efron, 2008).

In this study we advance a software called MultiPy, which is a Python-based open-source and freely-available toolkit for multiple hypothesis testing. Python has become in the recent past the programming language of choice for many scientists across several disciplines. Accordingly, there already exists general-purpose packages for scientific computing (van der Walt et al, 2011), data visualization and manipulation (McKinney, 2010), and machine learning (Pedregosa et al, 2011) to name a few. In neuroimaging, specialized Python software has been developed for example for the preprocessing, analysis, and source reconstruction of EEG and MEG data (Gramfort et al, 2013; 2014). While these and other packages allow efficient analysis of single-subject neurophysiological and neuroimaging data, most provide only a limited number of options for correcting group-level results for multiple comparisons. Thus, our aim was to develop software that fills this gap.

Taken together, we present here a novel model-based approach for comparing multiple hypothesis testing methods and quantify with simulated primary and follow-up experiments how large effect and sample sizes are needed to detect true effects reproducibly.

## 2. Methods

### 2.1. Definitions

In statistical null hypothesis significance testing (NHST), there are four possible mutually exclusive outcomes as summarized in Table 1. Specifically, there are two desired outcomes, which are true positives and true negatives, and two undesired outcomes, which are false positives and false negatives. We will focus here on the number of incurred false positives **V** while testing *m* hypotheses simultaneously under a specified critical level *α*, and how this count can be controlled using various procedures. We use the variables defined in Table 1 to denote the outcomes of NHST throughout the manuscript. Bold upper-case letters are used to denote random variables. Parts of the text also refer to *π*_0_, which is the proportion of true null hypotheses *m*_0_ among *m* tests, and also to 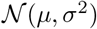, which is the normal distribution with location and scale parameters *μ* and *σ*^2^ respectively. The variable 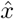 denotes an estimate of the variable *x*.

FWER is the probability of making one or more false positive conclusions while testing *m* hypotheses simultaneously. For independent tests, it can be described mathematically with the equation 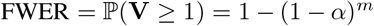 which is visualized in Figure 1A for the three conventionally used critical levels 0.001, 0.01, and 0.05. In practice, it can often be advantageous to control the FDR instead (Benjamini & Hochberg, 1995), which is the expected proportion of discoveries that are false, since it allows exchanging a small number of false positives to an increased power.

**Figure 1:**
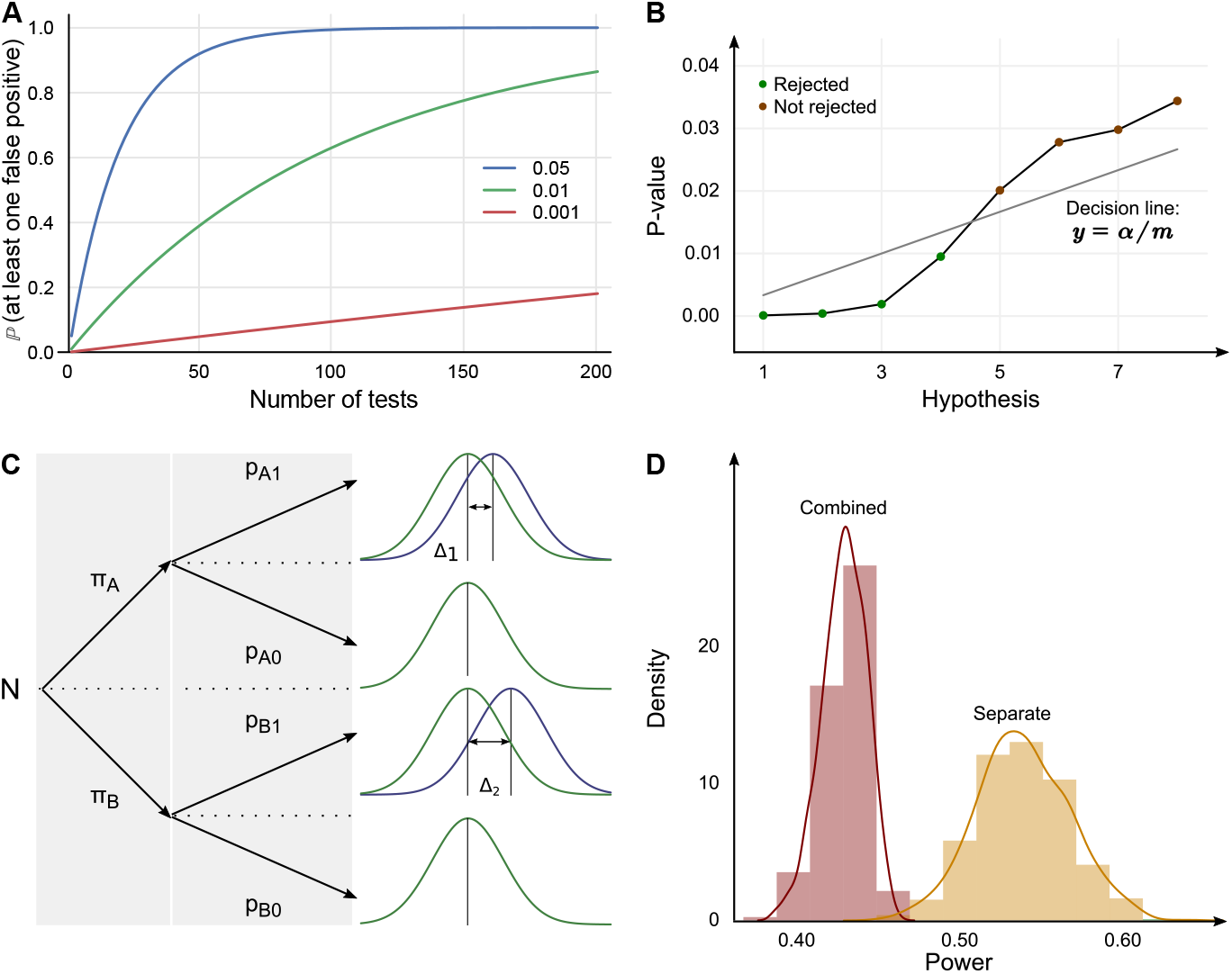
(A) False positives start to occur quickly when more than a few dozens of independent tests are performed simultaneously and no adjustment for multiple testing is made. The three lines correspond to the three conventional critical levels indicated in the figure legend. (B) The Benjamini-Hochberg FDR procedure can be interpreted graphically as rejecting sorted p-values that fall below the line *y* = *α*/*m* + 0 until the first non-significant p-value is seen. (C) Schematic of the structure of the separate-classes model. If information about the two separate classes were available a priori, performing two separate corrections would give a higher power than performing a single combined analysis (D). The histograms and density estimates were constructed by simulating data under the separate-classes model for 1000 iterations with the effect sizes Δ_1_ = 1.2 and Δ_2_ = 0.7.

### 2.2. Models for evaluating multiple hypothesis testing methods and performing numerical power and reproducibility analyses

#### 2.2.1. Spatial two-group model for comparing different FWER and FDR controlling methods

Fundamental effects in systems-level neuroscience data are often spatially or temporally continuous (Penny & Friston, 2003; Heller et al., 2006; Chumbley et al., 2010). To compare different methods for controlling the FWER and FDR under these circumstances, we first used the spatial two-group model suggested by Bennett and co-authors (2009). This model consists of a *n_v_* × *n_v_* variable two-dimensional grid with a *n_s_* × *n_s_* variable signal region in the middle. For each location on the grid, there are *N* samples in each of two groups denoted by A and B. The samples of the groups A and B are distributed as 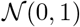 and 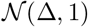 respectively in the signal region; the parameter Δ controls the effect size (Cohen’s *d*). In all other locations, the samples of both groups are distributed as 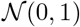 to model random noise. In other words, there is a true effect at every location within the signal region, and no true effects elsewhere. This model was used for the numerical comparison of the different multiple testing procedures, as well as performing prospective power and reproducibility analyses.

#### 2.2.2. Spatial separate-classes model for comparing different FWER and FDR methods when the true effects are distributed

In addition to being continuous in space, time, or frequency, fundamental effects in neuroscience data are also often distributed across temporal, spatial, or spectral scales (Heller et al., 2006; Chumbley et al., 2010). To compare the relative performance of multiple testing methods numerically when the true effects are distributed, we developed a separate-classes model which is a combination of two separate two-group models similar to Efron’s earlier study (2008) but with the spatial structure suggested by Bennett and co-authors (2009). In the model, there are now groups *A* and *B* in the first signal region and groups *C* and *D* in the second region. Similar to the two-group model, all samples outside the signal regions are distributed as 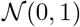 to model random noise. Within the signal regions, samples in groups *A* and *C* are distributed as 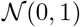, and samples in groups *B* and *D* as 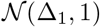 and 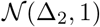 respectively. This model was used for testing the difference between performing two separate analyses and a single analysis, as well as testing how differences in the two effect sizes Δ_1_ and Δ_2_ influence the multiple testing results. The model is visualized schematically in Figure 1C.

### 2.3. Multiple hypothesis testing methods

#### 2.3.1. Classic methods for controlling FWER

Classic methods for controlling the FWER include the Bonferroni correction, Šidák’s correction (Šidák, 1967), Holm-Bonferroni procedure (Holm, 1979), and Hochberg’s procedure (Hochberg, 1988). The first two of these methods are similar with respect to computing new critical levels *α*_Bonferroni_ = *α/m* and *α*_Sidak_ = 1 – (1 – *α*)^1/*m*^ by a direct adjustment of the original level with the number of performed comparisons. In turn, the Holm-Bonferroni and Hochberg procedures apply the exact same threshold *α*/(*m* – *k* + 1) while correcting the *k*th ascendingly sorted p-value. The distinction between the two procedures is that they process the p-values in opposite orders and are hence categorized as step-down and step-up procedures.

#### 2.3.2. Benjamini-Hochberg FDR method

FDR controls the expected proportion of discoveries that are false, that is 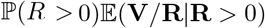 (Benjamini & Hochberg, 1995). Controlling the FDR implies willingness to accept a small fraction of false positives among the tests that are declared significant in exchange for improved power. The Benjamini-Hochberg FDR procedure is one of the most widely used FDR procedures, and can be understood graphically: p-values are sorted into ascending order and the hypotheses corresponding to the p-values that are below the line *y* = *α*/*m* + 0 are rejected until the first p-value crossing the line is seen (see Figure 1B).

#### 2.3.3. Adaptive FDR methods

The Benjamini-Hochberg FDR method assumes that *π*_0_ = 1 which can make it too conservative when the number of tested hypotheses is large (Storey & Tibshirani, 2003). Therefore, modern adaptive methods such as the Storey-Tibshirani q-value method (Storey & Tibshirani, 2003) and the two-stage procedure by Benjamini and colleagues (2006) attempt to estimate *π*_0_ as part of the correction procedure. While a successful estimation of *π*_0_ can allow greater power than the Benjamini-Hochberg method, there is a possible caveat: under some circumstances, applying the adaptive procedures may result in *more* significant discoveries than there were in the original uncorrected data (Reiss et al, 2012). In the study by Reiss and colleagues (2012), this kind of paradoxical results were found to occur in an MRI study with spatially widespread effects. The q-value method was found to be more vulnerable than the two-stage procedure (Reiss et al., 2012).

#### 2.3.4. Permutation testing

In comparison to the other approaches that control the FWER, permutation tests provide a non-parametric but more computationally expensive alternative. Instead of manipulating a set of p-values, they process the data under analysis directly. Permutation testing proceeds in two stages. In the first stage, the null hypothesis, exchangeability of observations under the null hypothesis, and the test statistic are specified (Nichols & Holmes, 2001). In the second stage, the permutation distribution is built by repeatedly relabeling the observations and computing the corresponding test statistics, which then allows calculating the significance of the correct labeling (Nichols & Holmes, 2001). Here, we performed the permutation tests using the procedure described by Maris and Oostenveld (2007) with a cluster-mass test statistic (Bullmore et al., 1999). However, in contrast to the original algorithm, we transformed the permutation test p-values into their upper bounds using the method suggested by Phipson and Smyth (2010) to avoid problems with zero p-values. In the original procedure described by Maris and Oostenveld (2007), obtaining zero p-values is possible when all permutation test statistics are less extreme than the observed one. The reason is that the p-value was defined as *p* = *b*/*m* where *b* is the number of times the permutation test statistic is equal or more extreme than the observed test statistic and *m* is the number of permutations. Therefore, the p-value could be underestimated by approximately 1/m when b is zero (Phipson & Smyth, 2010). However, this issue is avoided by defining the p-value instead as the upper bound *p_u_* = (*b* + 1)/(*m* + 1) which is always strictly positive (Phipson & Smyth, 2010).

#### 2.3.5. Random field theory

Random field theory (RFT) is an approach for controlling the FWER that has been widely used in the analysis of functional and structural MRI and positron emission tomography (PET) data (Worsley et al, 1992; Friston et al, 1994, Worsley et al, 1996), especially as part of the statistical parametric mapping (SPM) software (Frackowiak, 1997; Ashburner, 2012). Previous studies have also explored its applicability for the analysis of source-reconstructed EEG and MEG signals (Carbonell et al, 2004; Pantazis et al, 2005), and its strengths and weaknesses over permutation testing have been recently explored and discussed in depth (Eklund et al, 2016; Flanding & Friston, 2017). Application of RFT to neuroimaging data proceeds typically in three subsequent steps. First, the smoothness of the analyzed statistical map is estimated, which gives its resolution element or resel count. Here, we approximated the number of resels with the analyzed area divided by the squared full-width-at-half-maximum (FWHM) of the applied Gaussian smoothing kernel. Second, the expected Euler characteristic is computed for a range of thresholds, which can be informally defined as the number of distinct continuous regions that survive the thresholding. For two-dimensional data, the expected Euler characteristic is given by the equation 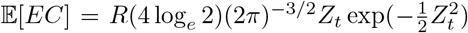 where *Z_t_* is the z-score threshold (Worsley et al., 1992). Finally, the empirical data is compared to the theoretical expectation for declaring statistically significant differences.

### 2.4. Data ana data analyses

#### 2.4.1. Comparison of multiple hypothesis testing methods using the spatial two-group and separate-classes models

The spatial two-group and separate-classes models were used to generate simulated data for numerically comparing the different FWER and FDR controlling procedures. The simulation using the two-group model was constructed as follows: the size of the signal region was 30 × 30 variables centered in the middle of a 90 × 90 variable grid containing no other true effects (i.e. the proportion of true effects was approximately 11%), and there were 25 samples in each of the two groups. The sample size was selected based on meta-analyses that suggest the median sample size in psychological and neuroscience research to be in the range 22–28 (Button et al, 2013; Szucs et al, 2017; Turner et al, 2018). The simulation was repeated for each effect size Δ in the range 0.5–1.5 with 0.05 increments. In the separate-classes model both signal regions were 15 × 15 variables and the overall grid size was 45 × 90 variables (i.e. the proportion of true effects was again approximately 11%), and the sample size was set at 25 in each group. The simulation was repeated with the effect size combinations {(Δ_1_, Δ_2_|Δ_1_, Δ_2_ ∈ [0.5 – 1.5]} with 0.05 increments. The critical level was set at α = 0.05 in both types of simulations. Also, the data was smoothed using a Gaussian kernel with FWHM set to match the size of the signal region when the RFT based method was applied. Further, all permutation tests were performed with 100 randomly drawn permutations with the t-threshold set at *t* = 1. Furthermore, if the spline-based null density estimator in the q-value method did not converge to the interval [0,1], the conservative choice of setting it as 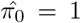 was made. Finally, the performance of each method was quantified as the number of rejected null hypotheses for which the alternative hypothesis was true (i.e. power).

#### 2.4.2. Reproducibility in the two-group model

To estimate effect sizes needed to observe true effects reproducibly in the two-group model, we performed simulated primary and follow-up experiments. The simulations were constructed as follows: the critical level was set at *α* = 0.05, the sample size at *N* = 25 in both of the two groups, the signal region was 30 × 30 variables in the middle of a 90 × 90 variable grid containing no other effects, the results were averaged over twenty realizations, and the tested effect sizes ranged from 0.5 to 1.5 with 0.05 increments. Tests that were declared significant in the primary experiment were selected for further testing in the follow-up experiment. To decide which hypotheses were reproducible across the simulated experiments, we used the FWER replicability method (Benjamini & Heller, 2008; Benjamini et al., 2009; Bogomolov & Heller, 2013). Briefly, this method defines weights *c_p_* and *c_f_* for the primary and follow-up studies respectively, with the constraint *c_p_* + *c_f_* = 1, and performs the multiple testing corrections at the critical levels *c_p_α* and *c_f_α*. Those tests that are declared significant in both the primary and the follow-up experiment are considered to be replicable and the rest are not. The simulations were performed by emphasizing the primary study at values 0.02, 0.5, and 0.98, which correspond to the critical levels 0.001, 0.25, and 0.049. Here, the selection of the lowest tested emphasis value was further motivated by a close correspondence to a recent suggestion of starting to user 0.005 as the new standard critical level in primary neuroscience experiments (Benjamin et al., 2018). The simulated data were analyzed using the correction methods introduced previously. The performance of each method was quantified with a reproducibility rate that is defined as *r* = *m_r_*/*m*_1_ where *m_r_* is the number of true effects declared reproducible and *m*_1_ = *m* – *m*_0_ is the number of all true effects.

### 2.5. Software implementation

MultiPy has been written in the programming language Python from the Python Software Foundation (http://www.python.org) (version 2.7.14). The implementation partly builds on previously published Python software packages, including Scikit-image (van der Walt et al., 2014) (version 0.13.0), SciPy (Oliphant, 2007) (0.17.0), NumPy (van der Walt et al., 2011) (1.10.2), Seaborn (0.8.0), and Matplotlib (Hunter, 2007) (2.1.0). The software is documented and available in http://github.com/puolival/multipy, and published under a permissive open-source license to facilitate further development and integration to other software and data analysis pipelines.

## 3. Results

### 3.1. Permutation testing is the most powerful method for analyzing data generated using the two-group and separate-classes models

The two-group and separate-classes models were used to simulate data at different sample and effect sizes to compare the relative performance of different multiple testing methods. The simulations were performed for a number of times, to evaluate the average power of each compared method at each sample and effect size. Permutation testing was the most powerful method for analyzing data generated using both the two-group model and the separate-classes model. Methods based on controlling the FDR produced intermediate results, and the least number of true positive effects could be detected using the other techniques that control the FWER. The differences in power between the different methods varied as a function of effect size. These results are visualized in Figures 2 and 3.

**Figure 2:**
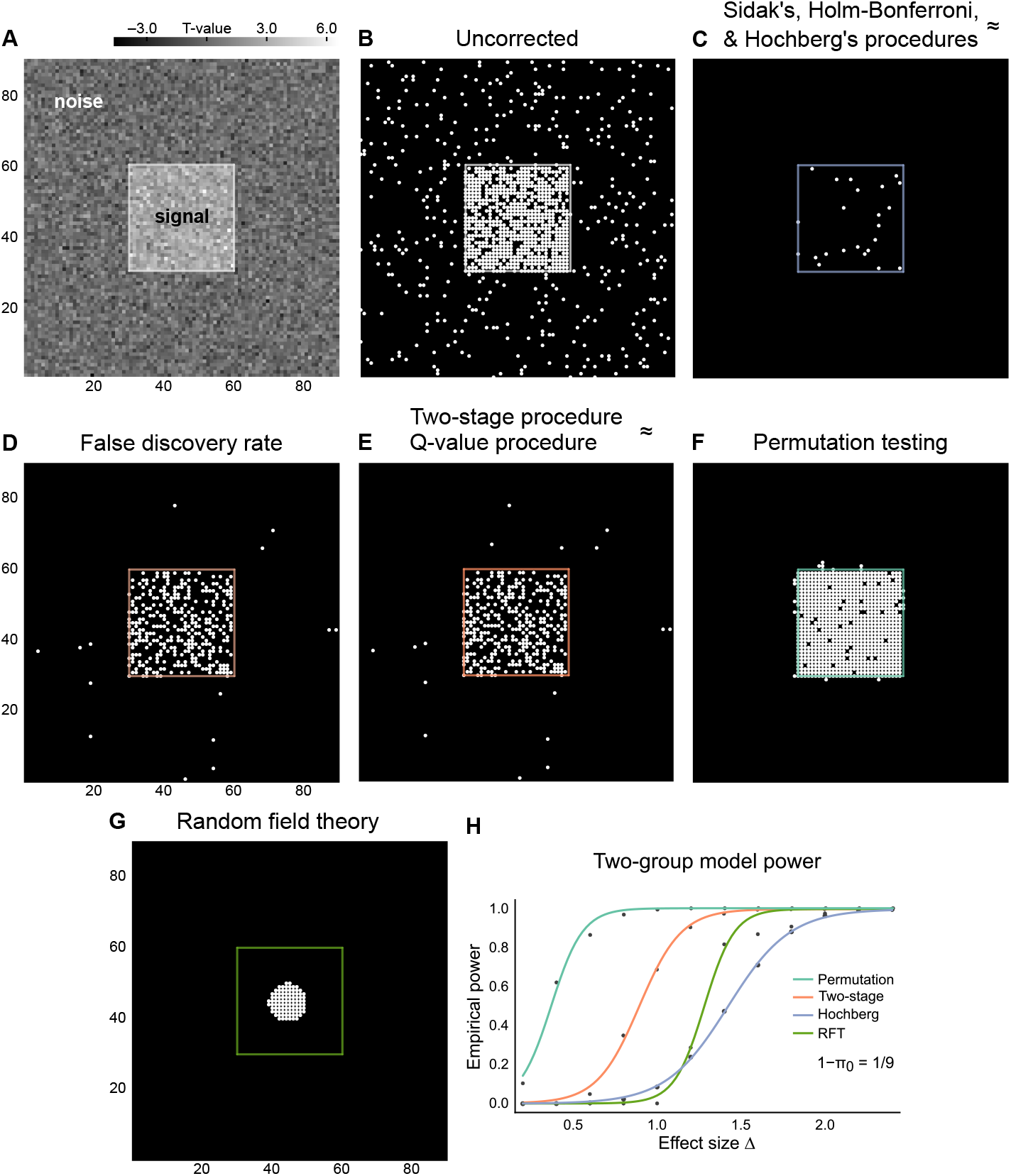
A single realization of the simulation performed using the two-group model. In each panel, there is a true effect at every location within the region outlined with green color and no true effects elsewhere. Repeating the simulation for a large number of times using various parameter selections allows estimating the relative merits of the different testing procedures, as well as performing power and reproducibility analyses. (A) The t-statistic of each variable. (B) The corresponding uncorrected p-values thresholded at the critical level *α* = 0.05. (C–G) The significant p-values after the correction for multiple testing has been performed. In this particular instance, permutation testing detects the largest number of true effects while incurring a small number of false positives. (H) The average empirical power for a representative selection of the different methods at various effect sizes when the simulation is repeated for several times.

**Figure 3:**
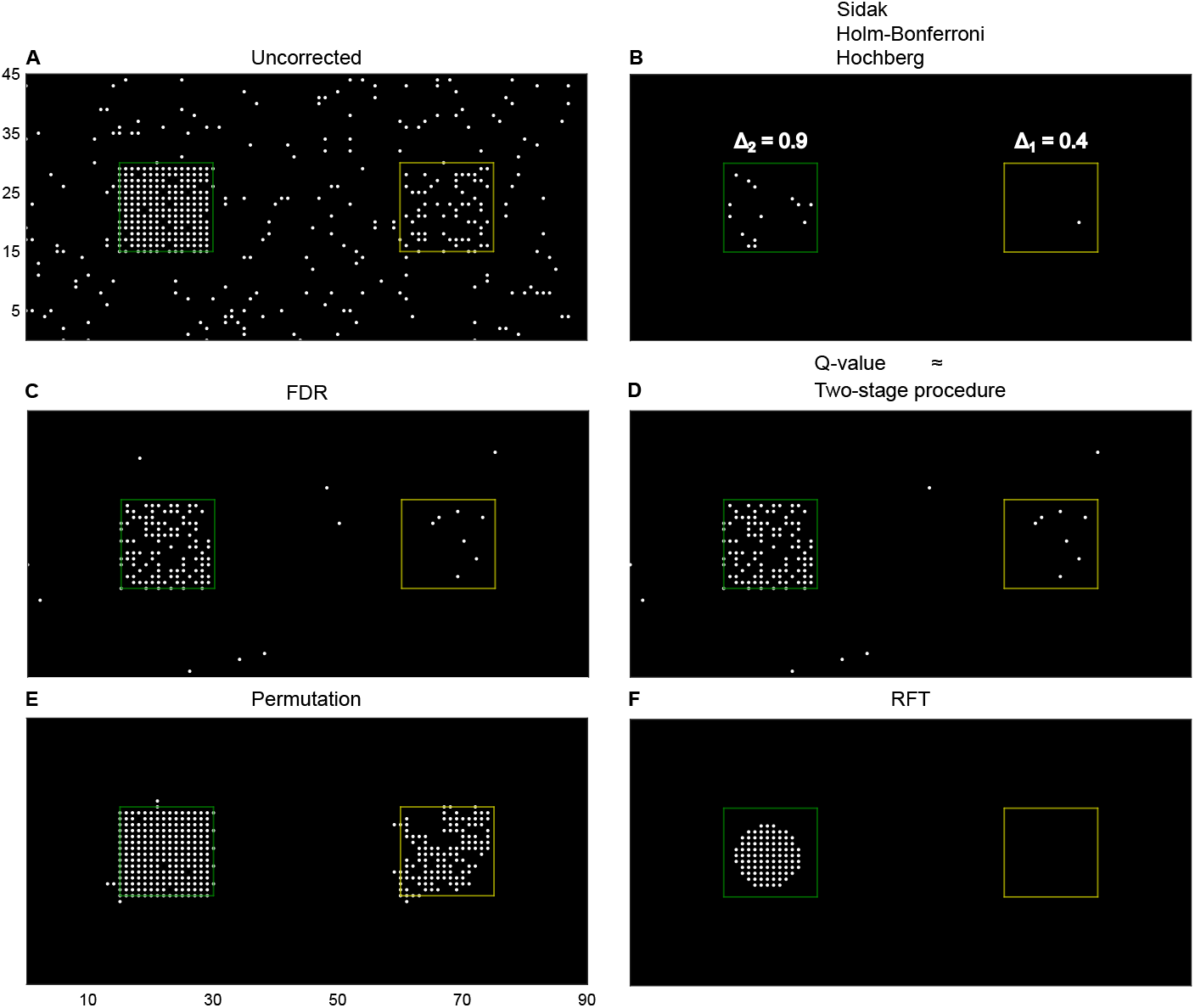
A single realization of a simulation performed using the spatial separate-classes model. There is a true effect at every location within the green and yellow regions and no true effects in the surrounding space. (A) The uncorrected p-values thresholded at *α* = 0.05. (B) The classic methods that control the FWER produced identical results, and therefore they were collapsed into one panel. (C) P-values that were significant when FDR was controlled using the classic Benjamini-Hochberg procedure. (D) Similar to (B), the adaptive FDR methods gave almost identical results, and therefore the results were also collapsed into a single panel. (E–F) Permutation testing detects both true effects almost entirely whereas RFT detects only most of the larger effect. Generally, repeating the simulation allows estimating the relative merits of the different techniques in the presence of two distinct effects. In addition, it is possible to test which of the two options is better: to perform two separate or a single combined analysis.

### 3.2. Placing an identical emphasis on the primary and follow-up studies produces optimal results

To evaluate what is the optimal multiple testing strategy between a primary study and a follow-up study, we simulated data using the two-group model at different sample and effect sizes and varied the relative emphasis placed on the primary study as well as the used correction method. This process was repeated for a number of times to evaluate the rate of reproducibility of true effects. The relative performance of the partial conjunction and FWER replicability methods was dependent on the emphasis placed on the primary study in comparison to the follow-up study assuming a conservative correction for multiple testing was carried out. When the primary study received larger emphasis than the follow-up study, the partial conjunction method outperformed the FWER replicability method. In contrast, when the follow-up study received higher emphasis than the primary study, the situation was reversed so that now the FWER replicability method produced the better results. These differences varied in magnitude as a function of the effect size (Figure 4A). When the FDR was controlled, optimal results were obtained when the primary and follow-up studies had an equal emphasis. This result was obtained repeatedly even though parameters of the simulations were varied; the same was true for a range of null densities, as well as for different sample sizes. However, the magnitude of the difference was again dependent on the effect size. To further probe the consistency of the results, the analysis was repeated with different correction methods, yielding consistent outcomes. These results are visualized in Figures 4B-C and 5. Another result obtained using the same set of simulations is that sample sizes larger than those typically used in neuroscience studies were needed to observe true effects reproducibly. A total of 40–50 samples per group or condition (i.e. a total of 80–100 samples) was needed to obtain rate a rate of reproducibility higher than 80% when the primary and follow-up studies were emphasized equally or the primary study was emphasized more than the follow-up study.

**Figure 4:**
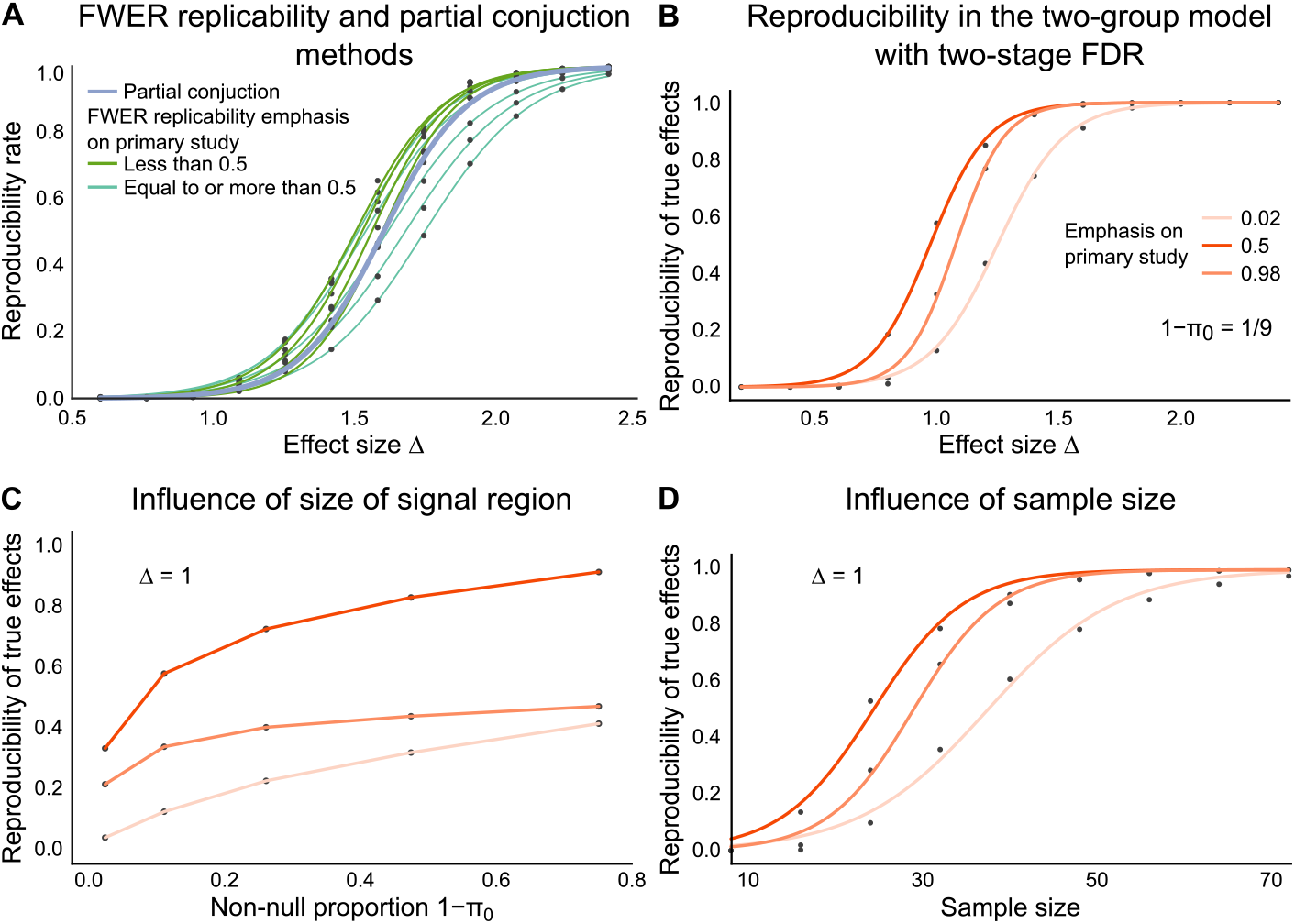
(A) A comparison of the partial conjunction and FWER replicability methods while performing a conservative multiple testing correction using the Hochberg’s method. The relative performance of the two methods depends on the emphasis of the primary study when analyzing the data using the FWER replicability method. (B) Primary and follow-up experiments were simulated using the spatial two-group model and the FWER replicability method was used to decide which hypotheses were reproducible across the two experiments. Now, the multiple testing correction was performed using the two-stage FDR procedure. The optimal result is obtained when the primary and follow-up studies are given equal importance, and not when strict corrections are performed in the primary study. (C) The result seen in panel (B) is not specific to a particular null density, or in other words, size of the signal region in the two-group model. (D) Sample sizes that are substantially larger than those typically employed in neuroscience experiments are needed to observe true effects reproducibly even at large effect sizes.

**Figure 5:**
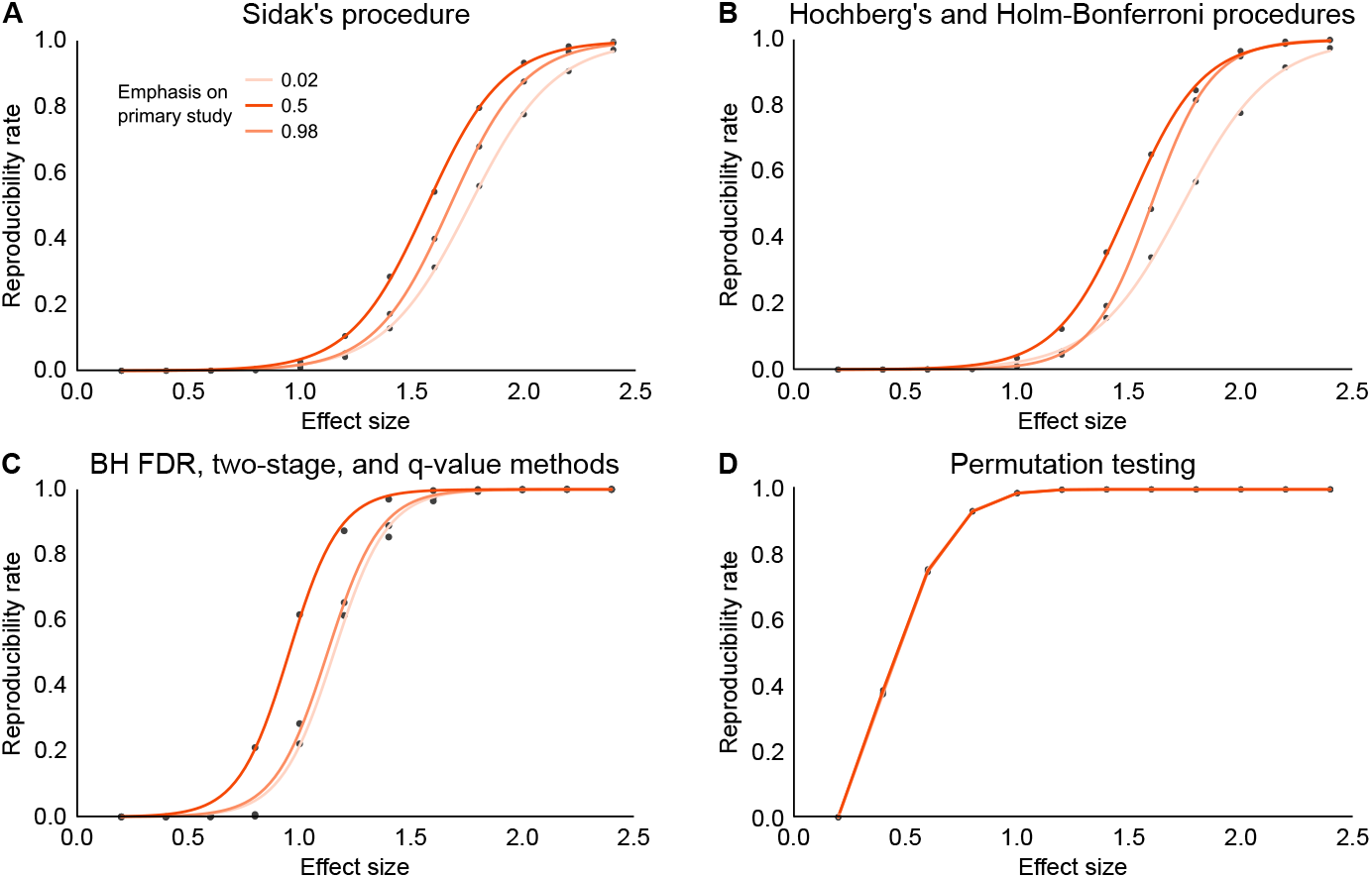
Placing an identical emphasis on the primary and follow-up studies produced an optimal number of reproducible true effects in the two-group model regardless of the used correction procedure (A–C). However, for permutation testing (D), this difference was negligibly small since there is only one cluster of true effects in the two-group model.

## 4. Discussion

We aimed here to quantify the performance of several multiple-hypothesis testing correction methods in the context of the reproducibility of the observations. To this end, we developed an open-source Python software, which implements classic and advanced techniques for controlling the FWER and FDR, simulating data under two different models, and performing reproducibility and power analyses. The software includes both parametric and non-parametric correction techniques, which were compared numerically using simulations performed under the two models. These models, namely the two-group model and the separate-classes model, allow capturing features typically observed in neurophysiological and neuroimaging data but are nevertheless as simple as possible. We obtained results that advance three points. First, the simulations showed that permutation testing is the most powerful approach for analyzing data generated under these models, with the caveat of having a high computational cost. Second, we found that incorporating prior knowledge to the testing process in the form of grouping hypotheses (i.e. performing separate analyses for distinct structures) can yield significant improvements in power. Third, we found that the combination of low power and testing of multiple hypotheses leads to poor reproducibility. This implies that sample sizes in neuroscience experiments may need to be substantially increased to enable a demonstration of true effects reproducibly. Moreover, these findings show that the recent suggestion of using 0.005 as the new threshold of statistical significance, instead of 0.05, in primary inferential statistics would not be optimal for reproducibility.

### 4.1. Permutation testing is the most powerful method for analyzing data generated using the spatial two-group model

For the data simulated using the two-group model, the classic methods that control the FWER (i.e. the Bonferroni, Šidák, Holm-Bonferroni, and Hochberg’s methods) produced identical or very similar results. In contrast, the Benjamini-Hochberg FDR procedure and the adaptive FDR methods detected expectedly more true positives while incurring a small fraction of false positives. Since the adaptive FDR procedures yielded similar results, it makes the two-stage procedure favorable over the q-value method due to its more stable null density estimator (Reiss et al, 2012). The permutation test outperformed all other approaches by a large margin, and the performance of the RFT based method was between the classic methods that control the FWER and the methods that control the FDR. Notably, while permutation testing was the most powerful approach here, it produced false positives that concentrated near the signal region boundaries. Indeed, one should be aware of its possible limitations in accurately establishing effect locations or latencies when applied to neuroimaging data (Sassenhagen & Draschkow, 2018). The power curves for representative methods for each class of correction methods are visualized in Figure 2H.

### 4.2. Incorporating prior knowledge to the multiple hypothesis testing process by grouping hypotheses of distinct structures increases power

The separate-classes model allows two signal regions with distinct effect sizes, which makes it possible to test whether prior information about the underlying data structure can be used to improve results obtained from the multiple testing process. Indeed, in the separate-classes model, detecting a second effect is more difficult if a single combined analysis is performed, in comparison to performing two separate analyses motivated by the available prior information. This results suggests that researchers should place more focus on considering whether their neurophysiological and neuroimaging data is most appropriately analyzed by combined or separate analyses. For example, it is well known that in EEG and MEG data the signal-to-noise (SNR) ratio decreases as a function of frequency, but yet most published frequency and time-frequency domain analyses were performed by combining all tests into a single analysis. We suggest future studies to leverage the available information about EEG and MEG measurement techniques, distinct roles of oscillations, components of evoked responses, trial structures, and other similar information while performing multiple testing corrections. Similar arguments can be put forward for functional and structural MRI data: consider for example the hemispheric lateralization of language, speech, and auditory functions.

### 4.3. Plan experiments for reproducibility

The recommended practice for sample size selection is to perform a priori power calculations. However, multiple testing has remained a consideration topic with no trivial solutions. In addition, reproducibility is typically not quantified in power calculations. Hence, it has remained largely unknown how well-powered studies are needed to observe true effects reproducibly. The issue is pressing since multiple testing occurs in most neurophysiological and neuroimaging research, and since the average study has an estimated power of only 8–31% (Button et al., 2013). Indeed, some scientific journals have already started to urge researchers to plan for reproducibility before conducting their experiments (Editorial, Nature Biomedical Engineering, 2018; Editorial, Nature Communications, 2018). To draw attention to this issue, we performed simulated primary and follow-up experiments using the spatial two-group model to highlight what the reproducibility rates might be at effect sizes typically observed in neuroscience research. Briefly, the results indicate that in this model the choice of multiple testing method and emphasis on the primary study have large influence on observing true effects reproducibly (Figure 4), and therefore we suggest similar analyses to be carried out while planning new primary and replication experiments. Overall, observing true effects reproducibly was difficult at small and moderate effect sizes with sample sizes that are typically used in psychological and neuroscience research. Our results also suggest that performing a too strict correction in the primary study is not optimal due to a substantially increased number of missed true effects, and that performing a too loose correction in the primary study is not optimal either, due to an increased number of false positive outcomes that must be subsequently verified at the replication stage. Instead, the optimal result was obtained by giving an equal emphasis for the primary and follow-up studies (Figure 4B); the same conclusion was reached using both the FWER replicability and the FDR r-value methods. Importantly, this finding is partly at odds with the recent suggestion of starting to use 0.005 as the new standard threshold of statistical significance in primary neuroimaging and neuroscience experiments as the solution to reproducibility problems (Benjamin et al., 2018). Indeed, the proposition does not take into account how data from the primary and follow-up experiments should be combined. Therefore, in contrast, we argue that researchers should first choose their desired critical level, and then perform prospective reproducibility analyses to plan their experiments.

### 4.4. MultiPy enables the evaluation of data analysis pipelines and assessment of reproducibility of planned experiments

Here we implemented an array of parametric and non-parametric multiple testing methods as well as two models for their numerical evaluation in a single open-source toolkit. Therefore, it is possible to use the provided software as a platform for developing new multiple testing methods, since their performance can be evaluated directly against other existing state-of-the-art techniques. Indeed, the two-group models have been extensively used for this purpose in the existing literature; see for example Heller & Rosset (2019) for recent work on optimal control of the FDR in such a model. The software can be also used to validate existing custom neuroimaging data analysis pipelines, which are presently abundant among the different laboratories (Carp, 2012), by simulating data at chosen effect and sample sizes and testing whether the FWER or FDR is controlled. Moreover, the software enables its users to perform prospective numerical power analyses, and importantly, evaluate the reproducibility of their planned experiments when the two-group or separate-classes models are good approximations to the effects seen in the empirical data. In addition, the developed software can be used in future studies as a platform for developing new multiple testing techniques, since it allows evaluating their performance directly against other existing state-of-the-art techniques. It also allows researchers to validate their custom data analysis pipelines using the two-group and separate-classes models; data can be simulated at chosen effect and sample sizes and then analyzed similar to real empirical data to test whether the FWER or FDR is controlled.

### 4.5. How to choose between controlling the FWER and FDR

A topic we have not discussed yet is how should one decide whether to control the FWER or the FDR? Generally, a strict control of false positives is often gained at the expense of false negatives, which may lead to important discoveries remaining unnoticed, especially when the number of tested hypotheses is large. Therefore, such tight control is best justified when any false positive discoveries are expensive due to ethical, financial, time, or other constraints. For example, one might wish to conduct a detailed confirmatory follow-up study corresponding to each discovery in the primary study; the use of animals or high research costs could make false leads too expensive. On the opposite side of the spectrum, a looser control is also a sound choice for many types of data analyses, since often there are no theories or models concerning each individual test. Here, consider for example a typical EEG or MEG induced-response study with two conditions and a comparison of the responses in the time-frequency domain as the main data analysis. The statistical testing would likely involve some hundreds or thousands of tests depending on the length of the analysis time-window and frequency resolution (or spacing of frequencies in the case of wavelet-based methods). In most cases like this, there would be no a priori expectations for all or even most of the time-frequency tuples. Instead, one would look for consistent activity patterns over several frequencies and time points; a small number of false positives within such regions would not compromise the results’ interpretation. Now, ideally, one would like to control the FWER but yet find a substantial amount of the true positive effects. This is the result that is obtained in our simulations with permutation testing, and therefore we suggest its use when the associated computational burden is not a major limitation and its assumptions are well met. Otherwise, the optimal choice between controlling the FWER or the FDR seems to mostly depend on the cost of false positives in the considered study.

### 4.6. Future directions

In the future, the approach advanced here can be expanded into several new directions. For example, topics that have been omitted here include multiple testing of multimodal (Winkler et al., 2016) and network data. Further, there are extensions to some of the implemented methods, such as FDR based RFT approaches (Chumbley et al., 2010) and covariate-adjusted FDR techniques (Genovese et al., 2006; Ignatiadis et al, 2016; Basu et al., 2018). Furthermore, there are many more specialized techniques such as those used for multiple testing of neuronal synchrony estimates (Maris et al., 2007; Singh & Phillips, 2010; Singh et al., 2011; Scott et al, 2015) that could be implemented as part of the software. In addition to these and other frequentist approaches, the software could be also developed to allow the use of Bayesian techniques, which are currently not widely supported by the major neuroscience software packages. The software can be also extended to include more data-generating models, which would help researchers to perform more diverse simulation-based power and reproducibility analyses. Finally, the software could be made directly compatible with data structures of existing Python-based neuroimaging analysis tools, which would facilitate its integration into existing data analysis pipelines.

## Acknowledgements

Authors thank Academy of Finland (grants 266745 and 281414) and Instrumentarium Science Foundation for funding, and Muriel Lobier, Sami Karadeniz, and Hamed Haque for helpful comments on an earlier version of the manuscript.

## Conflicts of interest

The authors declare no competing financial or other interests.

## Data availability statement

The data that support the findings of this study can be reproduced using the developed open-source software MultiPy, which is available online in https://github.com/puolival/multipy.

